# The inhibitory effect of a Corona virus spike protein fragment with ACE2

**DOI:** 10.1101/2020.06.03.132506

**Authors:** E. K. Peter, A. Schug

## Abstract

In this paper, we investigate the molecular assembly processes of a Coronavirus Spike protein fragment, the hexapeptide YKYRYL on the ACE2 receptor and its inhibitory effect on the aggregation and activation of the CoV-2 spike receptor protein at the same receptor protein. In agreement with an experimental study, we find a high affinity of the hexapeptide to the binding interface between the spike receptor protein and ACE2, which we investigate using 20 independent equilibrium MD simulations over a total of 1 *μ*s and a 200 ns enhanced MD simulation. We then evaluate the effect of the hexapeptide on the aggregation process of the spike receptor protein to ACE2 in long-time enhanced MD simulations. In that set of simulations, we find that the spike receptor protein does not bind to ACE2 with the binding motif shown in experiments, but it rotates due to an electrostatic repulsion and forms a hydrophobic interface with ACE2. Surprisingly, we observe that the hexapeptide binds to the spike receptor domain, which has the effect that this protein only weakly attaches to ACE2, so that the activation of the spike protein receptor might be inhibited in this case. Our results indicate that the hexapeptide might be a possible treatment option which prevents the viral activation through the inhibition of the interaction between ACE2 and the spike receptor protein.

**SIGNIFICANCE:** A novel coronavirus, CoV-19 and a later phenotype CoV-2 were identified as primary cause for a severe acute respiratory syndrome (SARS CoV-2). The spike (S) protein of CoV-2 is one target for the development of a vaccine to prevent the viral entry into human cells. The inhibition of the direct interaction between ACE2 and the S-protein could provides a suitable strategy to prevent the membrane fusion of CoV-2 and the viral entry into human cells. Using MD simulations, we investigate the assembly process of a Coronavirus Spike protein fragment, the hexapeptide YKYRYL on the ACE2 receptor and its inhibitzory effect on the aggregation and activation of the CoV-2 spike receptor protein at the same receptor protein.

## INTRODUCTION

In December 2019, a novel respiratory disease appeared in Wuhan, Hubei, China. Although it is still under debate, there are strong indications that a first cluster of infections occurred at the Huanan seafood market (1–3). A novel coronavirus, CoV-19 and a later phenotype CoV-2 were identified as primary cause for a severe acute respiratory syndrome (SARS) (4, 5). Within few days, the viral disease spread over whole China and within the following weeks, the local epidemic grew to a global pandemic with an exponentially growing infection rate. At present, the number of infected humans reached 3,855,788 with a number of 265,862 deaths associated with SARS-CoV-19 and CoV-2 (6). This global pandemic will have an unprecedented economic, sociological and political impact, in contrast to prior outbreaks of CoV related SARS epidemics (7). While a huge number of trials are still ongoing to develop a successful vaccination strategy against CoV-2, a direct medication of infected patients can have the potential to save lives and to stabilize the situation. The spike (S) protein of CoV-2 is the major target for the development of a vaccine or a potential strategy to tackle the viral entry into human cells (8–10). The S-protein forms trimers at the protrusions of the virus and comprises two functional subunits: S1 and S2. In the cascade of the viral entry, The S1 unit of the spike (S) protein facilitates the attachment of the virus at the surface of the cell (11). The S2 subunit, responsible for membrane fusion, employs TMPRSS2 for the S protein priming, while it uses ACE2 as entry receptor for membrane fusion (12–17). One of the key factors for its infectious potential for humans is the high conservation of ACE2 in different mammalian organisms (18), which allows its transmission from animals to humans. The receptor binding domain (RBD) of the S2 subunit contains five antiparallel beta strands, while alpha-helical and loop motifs form the connecting entities between the beta sheets. Between two beta sheets, an extended insertion forms the receptor binding motif (RBM), which binds to ACE2 at its N-terminal helix (19–21). Among a large number of potential targets, the inhibition of the direct interaction between ACE2 and the S-protein provides a suitable strategy to prevent the membrane fusion of CoV-2 and the viral entry into human cells (22, 23). In a combined experimental and theoretical study, a hexapeptide ((438)YKYRYL(443)) of the receptor domain of the SARS-CoV S-protein has been identified as an efficient inhibitor of the interaction between the S-protein and ACE2 (24). In vitro infection of Vero E6 cells by SARS coronavirus (SARS-CoV) was blocked by the hexapeptide. It also has been shown that the peptide inhibits the proliferation of CoV-NL63. Interestingly, the fragment (438)YKYRYL(443) carries the dominant binding epitope and binds to ACE2 with a high affinity of K(D)=46 *μ*M. Its binding mode was further characterized by saturation transfer difference (STD), NMR spectroscopy and Molecular Dynamics (MD) simulations. Based on this information the peptide can be used as lead structure to design potential entry inhibitors against SARS-CoV and related viruses.

In this article, we investigate the interaction of the hexapeptide (438)YKYRYL(443)) of the receptor domain of the SARS-CoV S-protein with ACE2 by MD simulations. MD has become a powerful *in silico* tool to complement experimental studies(25–28). We quantify the protein affinity to the binding site shown by Struck *et al*. (24). Second, we applied enhanced correlation guided MD simulations to measure the free energy of adsorption of the ligand to its binding site at ACE2. In the third stage of the study, we investigated the effect of the hexapeptide on the interaction of the S-protein with ACE2. In our simulations, we observe that the hexapeptide binds to the N-terminal region of ACE2 with a high affinity to 3 clusters that are located at the interface at which the spike receptor protein binds to the receptor as revealed in X-ray structures (19, 20). In the enhanced MD simulations, we observe that the spike receptor protein relaxes into an energy minimum which differs fundamentally from the X-ray structure: We find that the spike receptor protein rotates and binds to ACE2 at the N-terminal region by a hydrophobic patch, which is between residues Thr351 and Leu535. Our simulations suggest that the energetic minimum does not favour a hydrophilic interaction as shown in X-ray crystallography. In the simulations of binding of the S-protein receptor to ACE2 in the presence of the hexapeptide, the hexapeptide binds preferentially to the S-protein receptor in the vicinity to the ACE2 binding segment. Surprisingly, we find that this aggregation of the hexapeptide changes the aggregation process of the S-protein receptor, such that the activation of ACE2 is inhibited by the hexapeptide. Our simulations are in agreement with the experimental study and demonstrate the potential of the hexapeptide YKYRYL as a possible ‘new modality’ treatment option (29) which prevents the viral entry into human cells. Due to a damping effect by the cleavage of the peptide by proteases, a chemical modification which hinders that cellular process might increase the therapeutic potential of this peptide (29–31).

## METHODS

### Correlation guided dynamics (CORE-MD)

CORE-MD uses the path over the reduced action *L* as function of the momenta *p* and coordinates *q* along the trajectory (32–35):

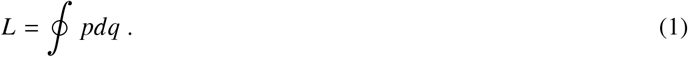

A path-dependent correlation function *C*(*t*) is defined:

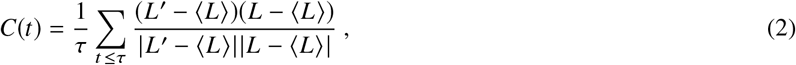

where 〈…〉 denotes the time average, where *L′* is determined at a time *t′* with a probability *p*:

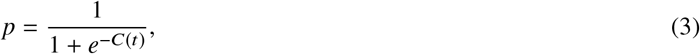

at every timestep. As a crucial element of the method, the correlation dependent density *ρ* as function of the correlation function *C*(*t*):

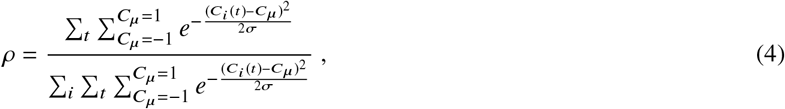

where *σ* defines the width of the Gaussian function (Due to the fact that we apply a histogram over 10^2^ bins, we apply *σ* = 2 × 10^-2^). Subsequently, we introduce a log pseudo likelihood function *l* of the correlation dependent density:

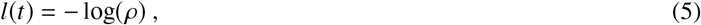

which defines the corresponding bias *A* with an additional parameter *α* with the units of an energy:

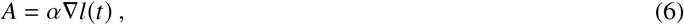

as the derivative along a unit vector with the length equal 1, due to the dimensionality of the correlation function. We scale the resulting gradient by a correlation dependent factor *r*, in order to enhance the decay of the auto-correlation and to achieve a faster access of the folded conformation space (For the proof, see supplementary information, section: I.A.). We define the factor *r* as:

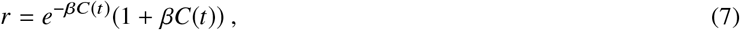

where *β* stands for a second constant.

### Simulation parameters and system setup

We centered the PDB structure of ACE2 (PDB: 6M0J, chain A) in a triclinic box with dimensions 7.419 x 8.361 x 8.614 nm^3^ fill the box with 17.026 SPC/E waters. For the preparation of 20 independent simulations over 50 ns, we placed the hexapeptide YKYRYL at 20 different initial positions in the vicinity to the potential binding site of ACE2 (see Figure 1 b). For the enhanced sampling simulation, we used one conformation to and simulated the system over 200 ns using the parameters *α* =5.0 and *β* = 0.5. In a third set of simulations we modeled a separated complex between ACE2 and the S-protein with an increased contact distance of approximately 2.2 nm by which the two domains are separated from each other. We modeled the separated complex with and without the hexapeptide. Both systems were centered in a triclinic box with dimensions 7.41900 × 9.13930 × 16.23050 *nm*^3^ (see Figure 1 c, d). We use the AMBER99SB forcefield to describe the interactions (37). For the 20 independent MD simulations over a total of 1 *μ*s, we used the stochastic velocity rescaling algorithm in combination with the berendsen barostat to simulate the system at NPT conditions at 300 K and 1 bar using a timestep of 1 fs (38). The enhanced sampling simulations of the hexapeptide-ACE2 complex and the two separated S-protein ACE2 complex simulations with and without the hexapeptide have been performed in implicit solvent using the standard GBSA AMBER99SB parameters. We measured the affinity to a specific binding site using the number of counts *N* in which the hexapeptide resides below a contact threshold of the *Cα* atoms of 0.65 nm in relation to the total number of frames in the trajectory *N_t_*. We define the relative affinity by the fraction of the affinity *η* divided by the maximal affinity *η_max_* measured for the specific system. The total affinity *ϵ* is given by the logarithm of the relative affinity:

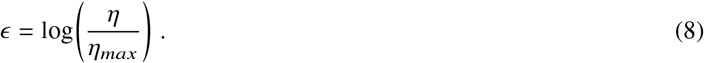

**Figure 1:**
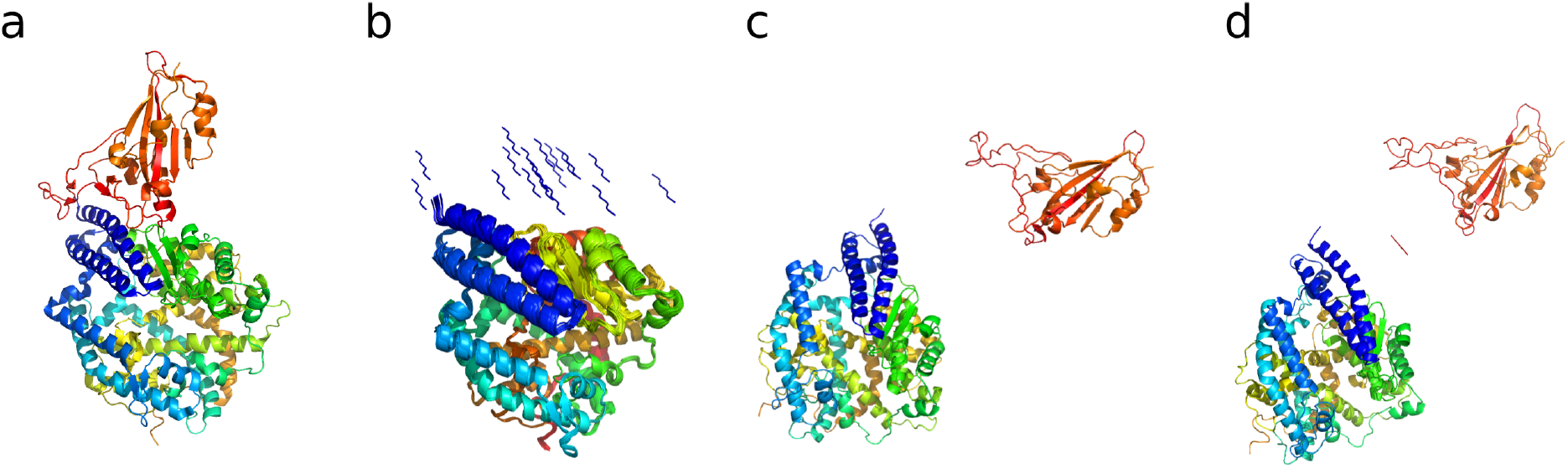
(a) Crystal structure of the SARS CoV-2 spike receptor domain bound with ACE2 (PDB: 6M0J (19)). ACE2 shown in blue, cyan, yellow and green color, the Spike receptor domain is shown in orange. (b) 20 different hexapeptide (YKYRYL) conformations used as initial starting models of independent 50 ns equilibrium MD simulations in explicit solvent. (c) Starting structure of the enhanced aggregation simulation of the spike receptor domain to ACE2. (d) Starting structure of the enhanced aggregation simulation of the same aggregate in the presence of the hexapeptide.

We define the free energy Δ*F* by the propability *P* along 2 order parameters (i.e. the distances *d*_1_ and *d*_2_ between pairs of atoms):

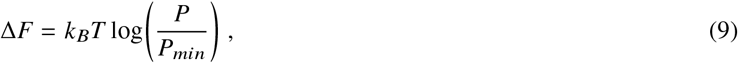

where *P_min_* stands for the minimal probability by which the histogram is populated. We used the GROMACS version 4.6 package for the equilibrium MD simulations and a modified GROMACS version 4.5.5 for the enhanced correlation guided simulations (39). We identified the preferential binding site of the hexapeptide at ACE2 through a distance based clustering. We then used the conformation of the hexapeptide at ACE2 for the calculation of protein interaction energies using the PRODIGY program (36). We modeled the individual hexapeptide conformers using the PyMOL modeling program (40).

## RESULTS AND DISCUSSION

### Simulations of hexapeptide assembly at ACE2

We tested the affinity of the hexapeptide to the ACE2 protein in 20 independent equilibrium MD simulations over a total simulation time of 50 ns. For this first set of simulations, we modeled 20 different starting conformations of the hexapeptide at an approximate distance of 1 nm away from the potential binding site between ACE2 and the S-protein. In order to examine the specific affinity of the hexapeptide for binding sites at ACE2, we determined the complete *Cα* – *Cα* distance matrix and averaged over all 20 trajectories. We find that the hexapeptide binds with 55 % of the total affinity to the interface between the N-terminal helix and a *β*-sheet located at Glu329 and a lowered contact propensity of 43-50 % to the residues Asn330 and Trp328 (see region (4) in Figure 2 a, c). We observe another contact cluster at Asp382 and Met383, where the affinity reaches values of 13 and 15 %. A third contact cluster is located at Thr55, where the affinity of the hexapeptide to ACE2 reaches a value of 21 % (see Figure 2 a, c).

**Figure 2:**
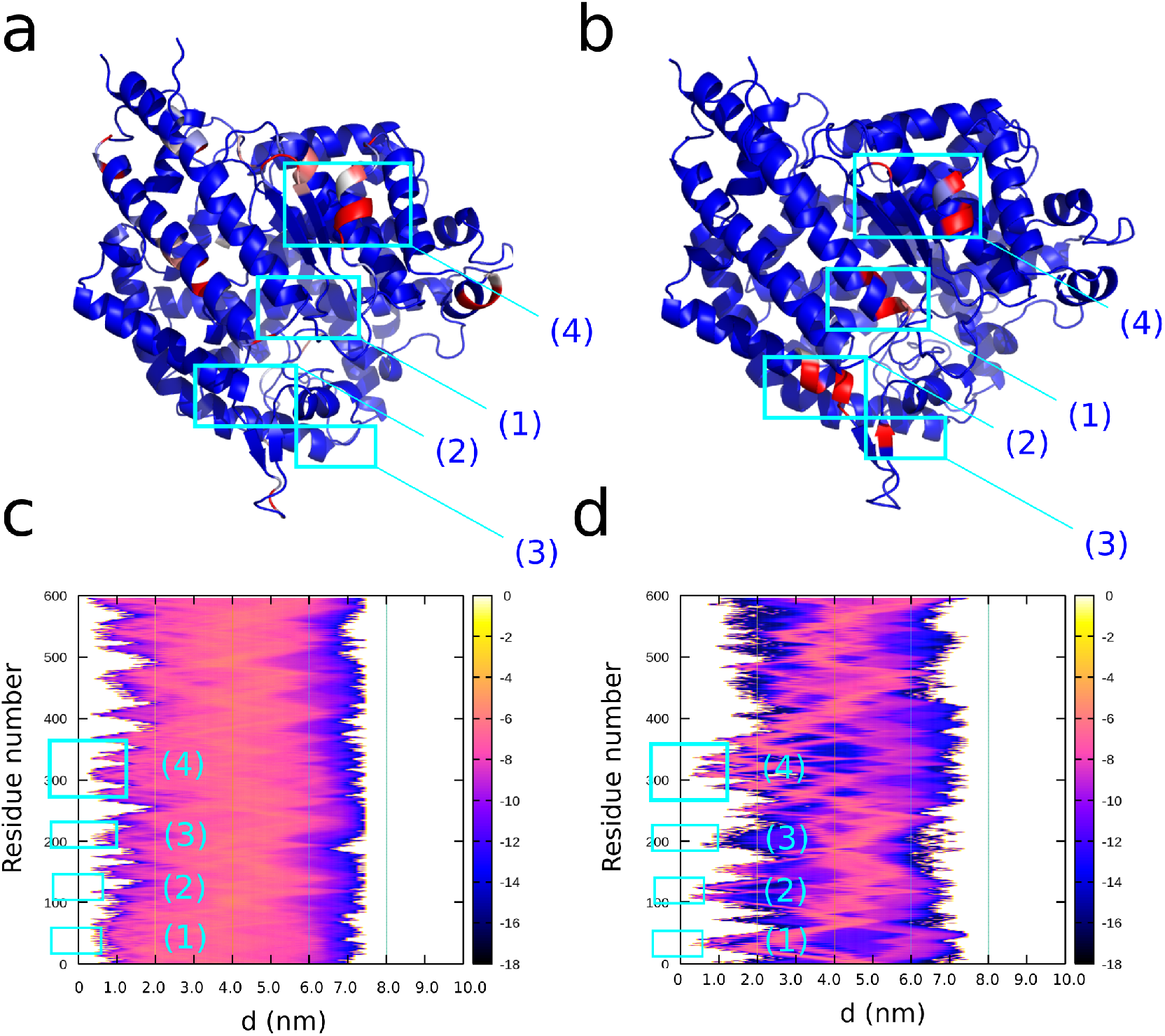
(a, b) Result from 20 equilibrium MD simulations over 50 ns of the hexapeptide in the vicinity of ACE2. (a) B-factor assigned structure of ACE2 with the affinity of the hexapeptide for regions at ACE2 indexed with a color gradient ranging from 0 (blue) to 100 % (red). (c) Log-plot of the relative affinity *ϵ* averaged over the set of 20 MD simulations. The affinity expresses the propensity of finding the hexapeptide as function of the residue number and the distance. (b, d) Result from the enhanced aggregation simulation over 200 ns of the hexapeptide in the vicinity of ACE2. (b) B-factor assigned structure of ACE2 with the affinity of the hexapeptide for regions at ACE2 indexed with a color gradient ranging from 0 (blue) to 100 % (red). (d) Log-plot of the relative affinity *ϵ* of finding the hexapeptide as function of the residue number and the distance averaged over the correlation guided MD simulation.

In an enhanced MD simulation, we simulated the assembly of the hexapeptide at ACE2 in implicit solvent. We used this simulation to cross validate our equilibrium MD simulations. In the enhanced MD simulation over 200 ns, we observe approximately identical binding patterns of the hexapeptide to the surface of ACE2. We observe a first cluster of contacts at Gly354 with an affinity of 27 %. We find a second contact cluster with a propensity ranging from 80 to 100 % is located at Trp328 and Glu329 (see region (4) in Figure 2 b, d). We observe another contact pattern at Gln325 with a propensity of 96 %. A fourth contact cluster is located at Leu132 with a relative propensity equal 56 %. Finally, we observe a last cluster of contacts at the N-terminus of ACE2 between the residues Ser124 and Gly130 with affinities ranging from 1 to 71 % (see regions (2) and (3) in Figure 2 b, d). An additional minor contact formation with a propensity of 2 % resides at Glu57 (see region (1) in Figure 2 b, d).

Our set of 20 equilibrium MD simulations agrees with the enhanced MD simulation on the general contact patterns of the hexapeptide at ACE2. In both sets of simulations, we find a major binding pattern in the vicinity of the N-terminal region of ACE2, where specifically Arg4 binds to Asn322, which is in agreement with the experimental study (24). The residues Tyr3 and Tyr5 are interacting with Ala387 in a hydrophobic binding mode (see Figure 3 a, b). We conclude that the hexapeptide binds preferentially to the N-terminal helix and the helical interface close to the N-terminus, which indeed blocks the binding interface between the spike receptor domain of CoV-2 and ACE2 (19, 20). Based on our findings, the hexapeptide shows a high affinity for the ACE2 binding region, which has the potential to inhibit the spike receptor protein activation, membrane fusion and the viral entry into the human cell (24). Subsequently, we used the conformation of the hexapeptide at ACE2 to model five further peptide variants and used an protein binding energy predictor (36) to determine the interaction strength of the models with ACE2 (see Figure 3 c, d). We find that the hexapeptide YKYRYL binds with −7.4 kcal/mol and *K_d_* = 3.4 × 10^-6^ M. A modification of the C-terminal residue to arginine and a hydrophobic modification at the position 4 to Leu leads to a lower interaction energy, as we find for the models YNYLYL and YNYLYR (Δ*G* =-6.8 and −7.1 kcal/mol). A mutation at the position two to Leu has only a moderate effect and leads to an interaction energy equal −7.3 kcal/mol. In contrast to the other variants, we found that Lys at position two should remain conserved, while a replacement of Arg at the position 4 with Asn increases the affinity of the hexapeptide to energies equal to −7.6 kcal/mol. The conformation with the lowest dissociation constant Kd is the variant YKYNYI, where the C-terminal Ile stabilizes the interaction leading to a value *K_d_* = 2.8 × 10^-6^ M.

**Figure 3:**
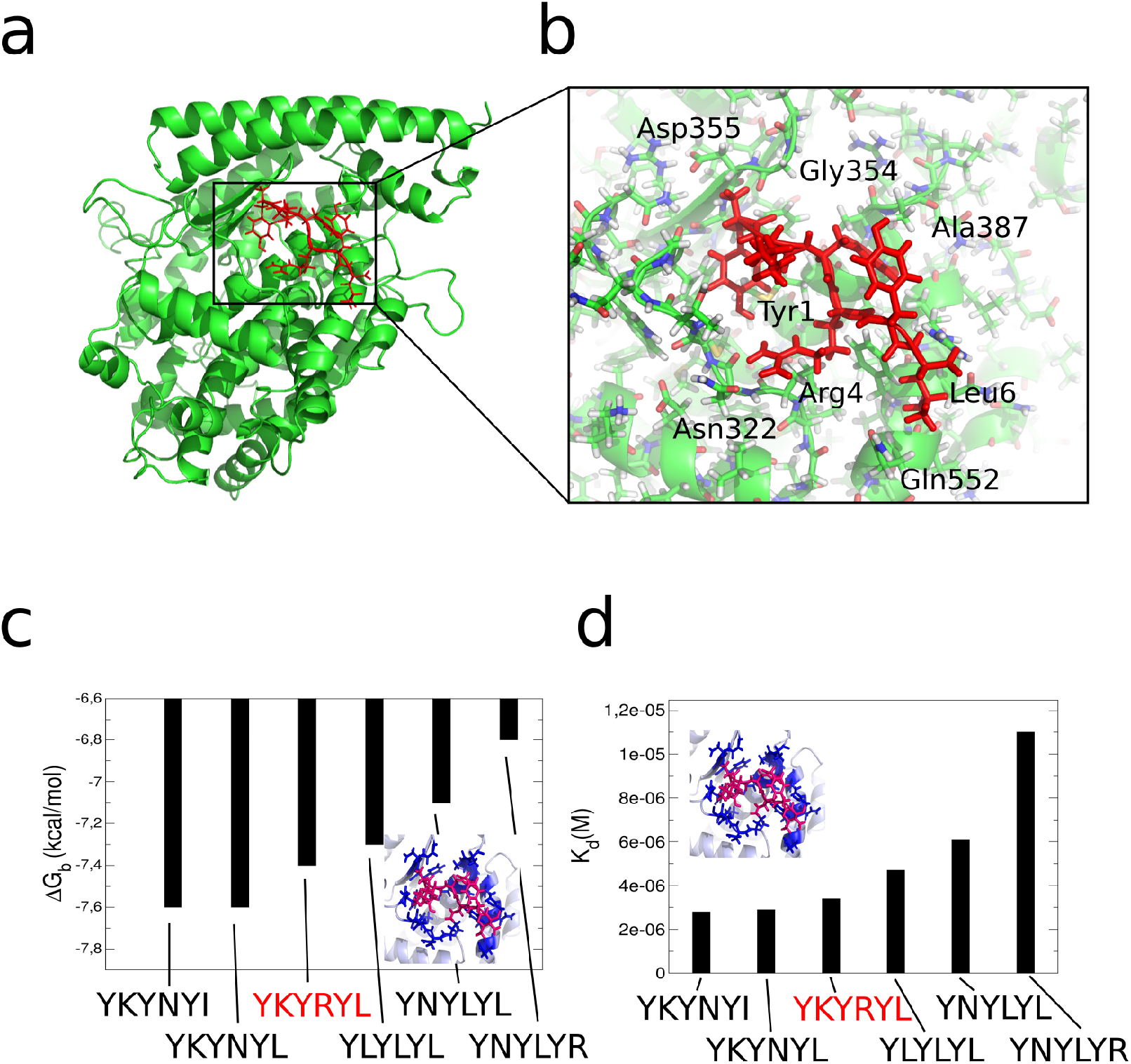
(a) Main conformation of the hexapeptide in complex with the N-terminal helical interface of ACE2 as revealed from equilibrium MD and enhanced sampling MD simulations. (b) Molecular view on the binding site of the hexapeptide at the interface with ACE2. (c) Results from protein binding energy calculations Δ*G* (36) on the hexapeptide YKYRYL and 5 further variant models in the binding site at ACE2. (d) Dissociation constant from the protein binding energy calculations on the 6 peptide variants.

### Simulations of spike receptor protein binding

We tested the effect of the hexapeptide on the assembly process of the spike receptor protein on ACE2 (see Figure 4 and 5). Therefor, we used a starting structure in which the spike receptor protein is separated by a distance of 2.2 nm away from the surface of ACE2 (see Figure 1 c, d). We simulated the system in and without the presence of the hexapeptide using correlation guided enhanced sampling. In the simulation of the assembly process of the spike receptor protein to ACE2, the hydrophilic interface formed by the loop region between the *β*-sheets of the spike receptor domain rotates away from the surface of ACE2, mainly due to a electrostatic repulsion. In a comparatively fast translatory process, the spike receptor domain binds to the N-terminal helix (blue) of ACE2 (see Figure 4 a, c). The interface between the spike receptor domain and ACE2 is mainly stabilized by hydrophobic interactions between residues in the range between Leu5354 and Thr351 (Spike receptor protein) and Gln60 to Glu75. Especially a hydrophobic interaction between Leu63 and His537 plays a major role in the stabilization of the spike receptor domain at ACE2 (see Figure 4 e). The relaxed final conformation of the spike receptor domain at ACE2 is rotated by approximately 90° in its attached orientation, when we compare the structure in an overlay with the experimental X-ray structure (PDB: 6M0J (19)) (see Figure 5 a). Additionally, the global contact pattern changed from a hydrophilic interface to a hydrophobic interaction at a different region of the spike receptor domain, which is initialized by a rotatory motion in the beginning of the simulation that is induced by an electrostatic driving force.

**Figure 4:**
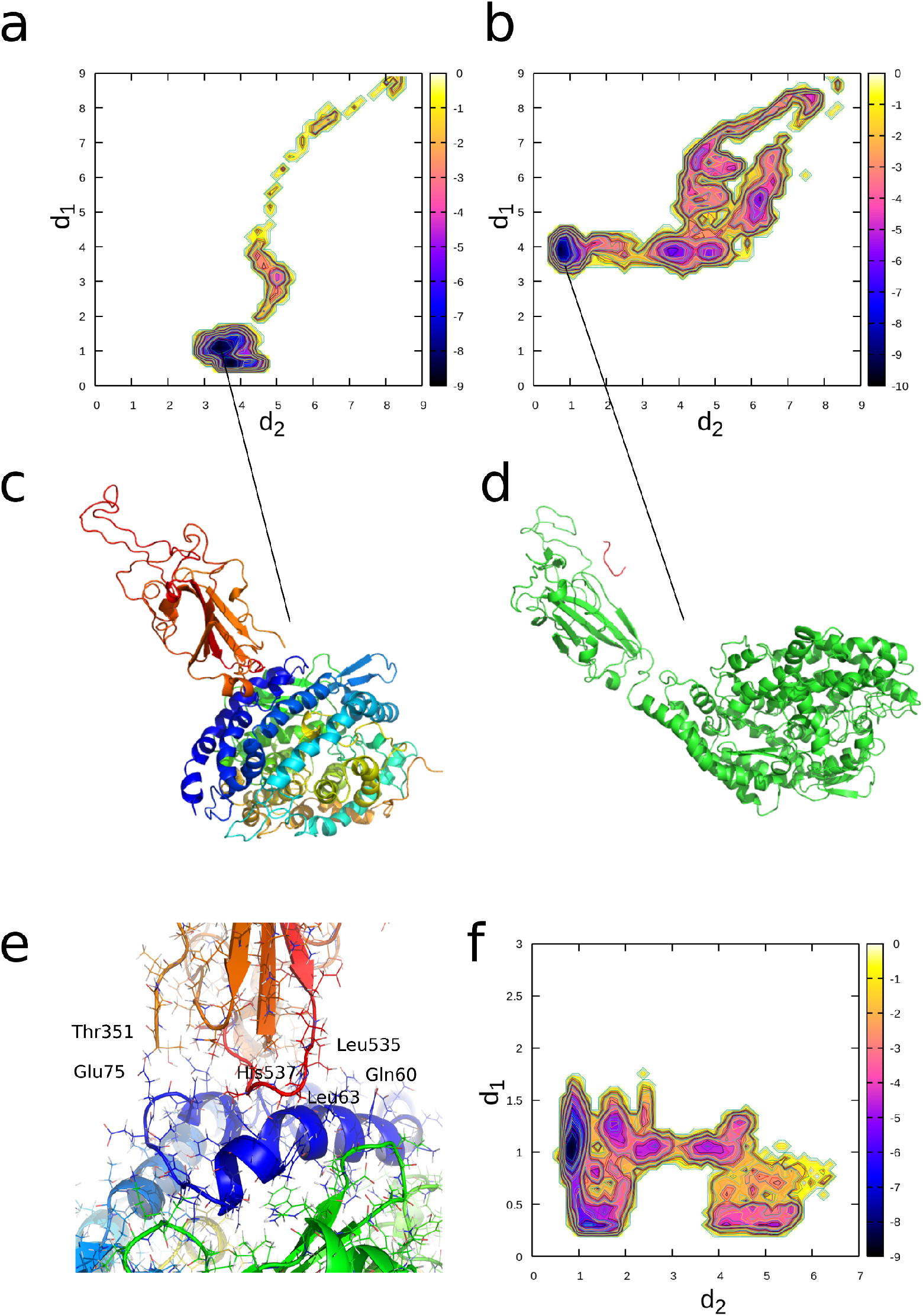
Results from enhanced sampling MD simulations of the aggregation process of the spike receptor protein with ACE2 in and without the presence of the hexapeptide. (a) Free energy landscape averaged over the trajectory of the spike-receptor protein - ACE2 aggregate without the hexapeptide as function of the order parameters *d*_1_ and *d*_2_, given by the distances between the residues Ile21 CB (ACE2) - Val524 CB (Spike-receptor) (*d*_1_) and Ala65 CB (ACE2) - Leu390 CB (Spike-receptor) (*d*_2_). (b) Free energy landscape averaged over the trajectory of the spike-receptor protein - ACE2 aggregate in the presence of the hexapeptide as function of the order parameters *d*_1_ and *d*_2_, given by the distances between the residues Ile21 CB (ACE2) - Val524 CB (Spike receptor) (*d*_1_) and Ala65 CB (ACE2) - Leu390 CB (Spike receptor) (*d*_2_). (c) Final converged state of the spike-receptor protein in complex with the N-terminus of ACE2 in the simulation without the hexapeptide. (d) Final converged state of the spike-receptor protein in the presence of the hexapeptide. (e) Molecular view of the interface of the spike-receptor protein at ACE2. (f) Free energy landscape as function of the order parameters *d*_1_ and *d*_2_, given by the distances between the residues Lys2 NZ (Hexapeptide) - Glu465 OE1 (Spike-receptor) (*d*_1_) and Lys2 NZ (Hexapeptide) - Lys335 (ACE2) (*d*_2_). The units in the color bar are given in *k_B_T*.

**Figure 5:**
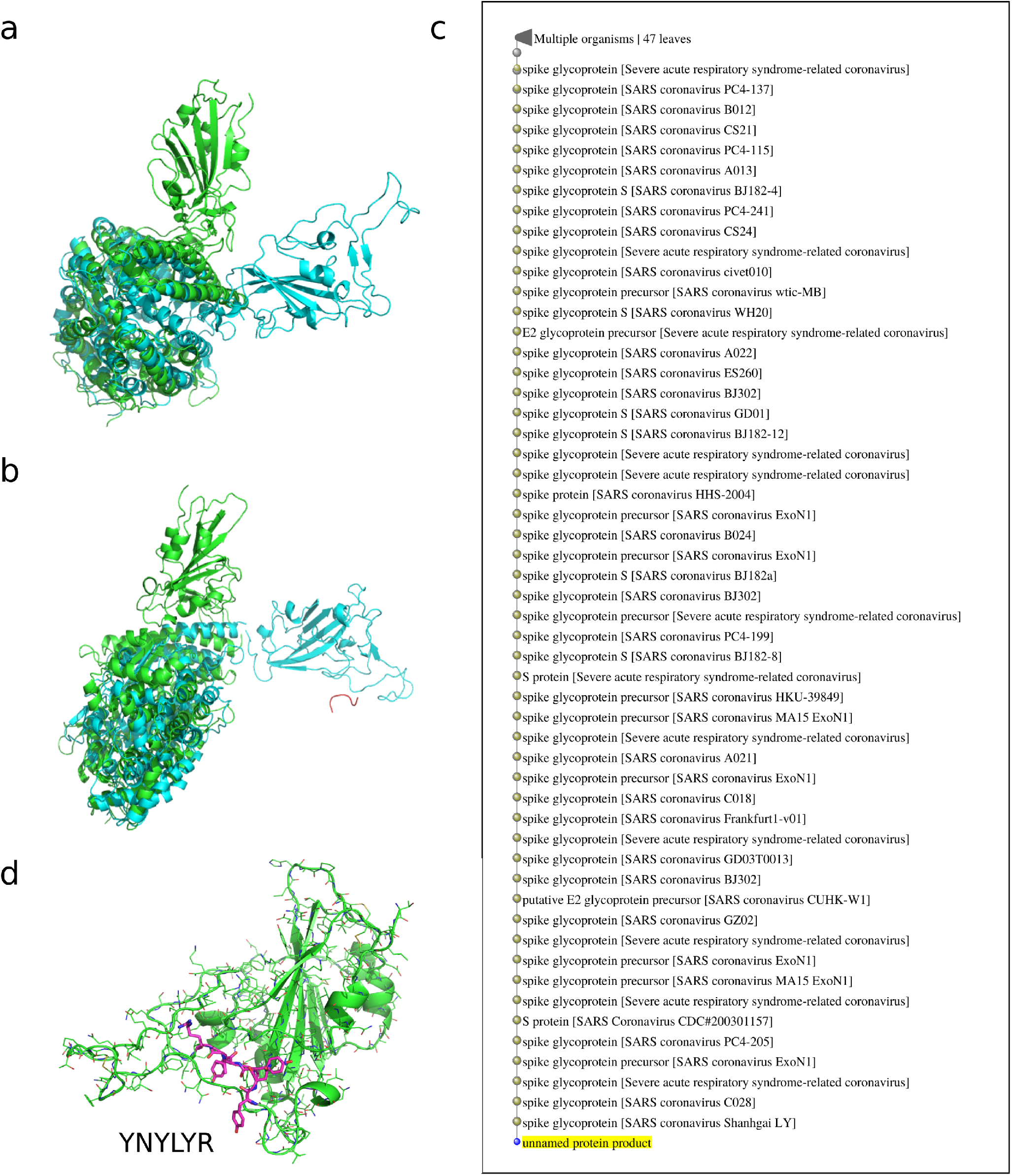
Structural overlays of the PDB structure (PDB: 6M0J) of the spike-receptor protein in complex with ACE2 (green) and the 2 final structures from the enhanced MD simulations without (a) and in the presence of the hexapeptide (b) (cyan). (c) List of SARS CoV viruses as a result from a BLAST search over the protein sequence space of all organisms. Surprisingly, the hexapeptide fragment preferentially occurs in SARS CoV viruses, which makes it suitable as potential drug, due to its dissimilarity with human proteins. (d) Hexapeptide fragment YNYLYR in the SARS CoV-2 spike protein receptor (PDB: 6M0J, chain E), indicating that the Tyrosine repeat at the positions 1,3 and 5 might be important for the design strategy of a potential peptide-mimetic for the treatment of SARS-CoV-2 infections.

In the simulation of spike receptor protein assembly to ACE2 in the presence of the hexapeptide, we observe a fast binding process of the hexapeptide to spike receptor protein at Glu465 (S-receptor protein) and Lys2 (Hexapeptide). We find a secondary contact between the hexapeptide and the spike-receptor domain between Ser349 (S-receptor protein) and Arg4 (hexapeptide) in the initial stage of the simulation. The hexapeptide then diffuses along the surface of the spike-receptor protein till its binds strongly to Glu465 (see Figure 4 d, f). An initial rotatory process of the spike receptor domain is highly analogous to the simulation without the hexapeptide, in which the binding motif of the spike-receptor protein rotates away from the surface of ACE2 due to an electrostatic driving force. In contrast to the simulation without the hexapeptide, we find that the spike receptor domain attaches only weakly at the peripheral region of the N-terminal helix 4 nm away from the conformation without the hexapeptide (see Figure 4 b, d and Figure 5 b). We anticipate that this conformation corresponds to an inhibited state, in which the spike-receptor does not become activated and the process of membrane fusion might get inhibited.

We were surprised by the high specificity with which the hexapeptide bound to ACE2 and the spike protein receptor. Since we observed that the hexapeptide inhibits the binding process of the spike protein receptor, we performed a BLAST search over all organisms, which contain the specific fragment in their protein sequences (41, 42). Surprisingly, we found that 47 out of 50 hits in the sequence search returned SARS Corona Virus organisms, while only 3 hits were contained in bacteria, which shows that the hexapeptide-pattern preferentially occurs in the SARS CoV viral organism, but not in human proteins (see Figure 5 c). That result shows that an *a priori* affinity for another function in the human organism can be excluded, which makes the hexapeptide a suitable candidate as a potential drug. When we analyzed the sequence of the SARS-CoV-2 corona spike receptor, we find a hexapeptide sequence YNYLYR, which contains the same Tyr repeat at the positions 1, 3 and 5, but different residues at the positions 2, 4 and 6, which might be an indicator that Tyr at the positions 1, 3 and 5 is imminent for the specificity of the spike receptor protein fragment (see Figure 5 d). However, we found that this hexapeptide sequence leads to the lowest interaction energy Δ*G*=-6.8 kcal/mol as we found in a modeling approach using the preferential hexapeptide ACE2 binding site as a structural model (see Figure 3 c, d). We only can speculate that the aminoacids at the positions 2, 4 and 6 are affecting the relative affinity of the fragment for the ACE2 receptor, while we find that Tyr at the positions 1, 3 and 5 is essential for the binding. We assume that Tyr at the positions 1, 3, and 5 has to be conserved for the design of a peptide mimetic used as potential drug against SARS CoV-2, while the hexapeptide sequence YKYRYL inhibits the viral interaction with ACE2 as we have shown in this work.

## CONCLUSIONS

In this paper, we investigated the assembly process of a Coronavirus Spike protein fragment, the hexapeptide YKYRYL on the ACE2 receptor and its effect on the aggregation and activation of the CoV-2 spike receptor protein at the same receptor protein. In agreement with an experimental study, we find a high affinity of the hexapeptide to the binding interface between the spike receptor protein and ACE2, which we investigated using 20 independent equilibrium MD simulations over a total of 1 *μ*s and a 200 ns enhanced MD simulation. We then evaluated the effect of the hexapeptide on the aggregation process of the spike receptor protein to ACE2 in long-time enhanced MD simulations. In that set of simulations, we found that the spike receptor protein does not bind to ACE2 with the binding motif shown in experiments, but it rotates due to an electrostatic repulsion and forms a hydrophobic interface with ACE2. Surprisingly, we observed that the hexapeptide binds to the spike receptor domain, which has the effect that this protein only weakly attaches to ACE2, so that the activation of the spike protein receptor might be inhibited in this case. Our results indicate that the hexapeptide might be a possible treatment option which prevents the viral activation through the inhibition of the interaction between ACE2 and the spike receptor protein.

## AUTHOR CONTRIBUTIONS

EP and AS designed the study. EP performed the simulations and analyzed the data. EP and AS wrote the manuscript.

## ACKNOWLEDGEMENT

We recognize support by the Impuls- und Vernetzungfond and an ERC recognition award of the Helmholtz Association. The authors gratefully acknowledge Jülich Supercomputing Centre support for COVID-19 research by providing computing time.

